# Largely unlinked gene sets targeted by selection for domestication syndrome phenotypes in maize and sorghum

**DOI:** 10.1101/184424

**Authors:** Xianjun Lai, Lang Yan, Yanli Lu, James C. Schnable

## Abstract

The domestication of diverse grain crops from wild grasses resulted from artificial selection for a suite of overlapping traits producing changes referred to in aggregate as ”domestication syndrome”. Parallel phenotypic change can be accomplished by either selection on orthologous genes, or selection on non-orthologous genes with parallel phenotypic effects. To determine how often artificial selection for domestication traits in the grasses targeted orthologous genes, we employed resequencing data from wild and domesticated accessions of Zea (maize) and Sorghum (sorghum). Many ”classic” domestication genes identified through QTL mapping in populations resulting from wild/domesticated crosses indeed show signatures of parallel selection in both maize and sorghum. However, the overall number of genes showing signatures of parallel selection in both species is not significantly different from that expected by chance. This suggests that, while a small number of genes will extremely large phenotypic effects have been targeted repeatedly by artificial selection during domestication, the optimization portion of domestication targeted small and largely non-overlapping subsets of all possible genes which could produce equivalent phenotypic alterations.

## Introduction

The characteristics of modern crops are the result of thousands of years of artificial selection applied consciously or unconsciously by farmers and plant breeders. An estimated 2,500 plant species have experienced some degree of artificial selection, with approximately 10% of these domesticated to the point that the species depends on humans for survival^1^. Of the hundreds of crops domesticated by human civilizations, three species – rice, wheat, and maize – provide more than half of all calories consumed around the world. These three crops all belong to the same family Poaceae (the grasses), a clade that has contributed a total of at least 48 domesticated crop species to human civilization, including at least 30 species domesticated as sources of grain^2^. Artificial selection for grain production produced a suite of shared phenotypic changes in grain crops referred to as ”the domestication syndrome”^3^. Grain crop domestication syndrome includes loss of seed shattering, increased apical dominance, more uniform maturity across inflorescences and across tillers, increase in size and/or number of inflorescences, larger seeds, greater carbohydrate content and lower protein content per seed, and reduction or loss of seed dormancy^3^.

Quantitative genetic studies can be used to identify large effect genes involved in producing the changes associated with the domestication syndrome in crop species which are interfertile with their wild progenitors. In maize, an estimated five loci have large enough effects on domestication traits to be mapped using conventional QTL analysis^4^. Of these loci, several have been mapped including *teosinte branched1, tb1* where the allele selected for significantly reduces the development of tillers^5^, *teosinte glume architecturel, tgal* where the allele selected for abolishes the stony fruitcase surrounding teosinte seeds^6^, and *grassy tillersl, gtl* where the allele selected for results in many fewer ears per plant during domestication^7,8^. In rice, many functionally characterized genes that underlie phenotypic changes during the domestication have also been identified through QTL mapping, such as the *Seed dormancy 4, Sdr4* where the allele selected for acts produces a reduction in seed dormancy^9^, *Tiller Angle Controll, TAC1* where the allele selected produces a reduction tiller angle producing more photosyntheticly efficient canopy architecture^10^, and *betaine aldehyde dehydrogenase2, BADH2* a loss of function allele selected for during domestication results in the accumulation of 2-acetyl-1-pyrroline in fragrant rice^11^.

A second approach to identify loci which were targets of artificial selection is to look at changes in the diversity and frequency of haplotypes at particular regions of the genome between populations of a crop species and populations of wild relatives. Notably, while quantitative genetic evaluation of recombinant populations generally identifies less than ten large effect loci responsible for many of the differences observed between domesticated grain crops and their wild relatives, genome wide population genetic approaches generally identify hundreds to thousands of loci as targets of selection during domestication in the same species. Hufford and co-workers used resequencing data from a set of 75 teosinte and maize lines to identify 484 regions of the maize genome likely to have experienced selection during transition from wild teosintes to maize landraces and another 695 regions likely to have experienced selection during transition from – largely tropical – landraces to – largely temperate – elite lines^12^. Huang and co-workers used genome resequencing data of 1,083 cultivated rice and 446 wild rice to identify 55 selection regions encompassed 2,547 candidate artificially selected genes during the domestication from wild to cultivated rice^13^. In sorghum, a set of 725 genes which were likely the targets of artificial selection during domestication and/or crop improvement were identified from resequencing of 44 lines of sorghum and wild relatives^14^.

Parallel phenotypic changes which are parts of the domestication syndrome in grain crops could result from parallel or lineage-specific changes at a molecular level. A recently published study demonstrated the the loss of seed shattering resulted from disruption of the same gene in maize, sorghum, and rice^15^. *Heading Date1* is a major QTL controlling flowering time which shows evidence of being under parallel artificial selection during the process of domestication for sorghum, setaria, and rice^16^. Flowering time QTL identified in a population of wild x domesticated *Setaria* lines co-localize with flowering time QTL identified at syntenic orthologous locations in the genomes of maize and sorghum^17^. A significant number of candidate genes associated with seed size exhibited signals of parallel selection during domestication in maize, rice, and sorghum^18^. However, not all parallel phenotypic changes produced by artificial selection result from parallel evolution at molecular level. Artificial selection for adaption to high altitudes in different maize populations targeted largely unrelated sets of genes in mexican and andean highland populations^19^.

Here we focus on two grain crops, maize (*Zea mays* ssp. mays) and sorghum (*Sorghum bicolor* ssp. *bicolor*, which diverged from a common ancestor approximately 12 million years ago^20^. Maize was domesticated from Balsas teosinte (*Zea mays* ssp *parviglumis*) in mesoamerica, with a center of origin in the lowlands of southwestern Mexico^21^. Sorghum is believed to have first been domesticated from broomcorn (*Sorghum bicolor* ssp. *verticilliflorum* in Ethiopia^22^, with a potential second independent domestication in west Africa^14,23^. Major QTLs for seed shattering, seed mass, and flowering time were localized to syntenic regions between sorghum and maize^24^. The parallel set of phenotypic changes during domestication in these two species^3^, the high degrees of conserved collinearity across grass genomes^25,26^ and the bias towards genes with detectable phenotypic effects being conserved at syntenic locations across grass genomes^27^, offer an opportunity to test the hypothesis that the parallel phenotypic changes in maize and sorghum resulted from artificial selection acting on orthologous genes in both species. We find that genes conserved at syntenic orthologous locations in maize and sorghum were significantly more likely to be targets of selection during domestication than non-syntenic genes unique to one species. In maize, domestication preferentially targeted genes on the dominant maize1 subgenome rather than their retained duplicates on the maize2 subgenome. Genes identified through quantitative genetic studies of domestication traits in one species were likely to show signatures of selection in the other species, however the overall overlap between genes identified using population genetic methods in both species was only marginally greater than expected by chance.

## Results

### Known Domestication Targets in Maize

A set of 16 genes shown to exhibit functional variation between maize and teosinte or between maize landraces and improved lines for traits linked to domestication based on single gene or single gene-family studies was assembled (Table 1). Changes to the gene *shattering1* are implicated in loss of seed shattering in maize^15^. Differences between the teosinte and maize alleles of *teosinte branched1* reduce the growth of axillary buds and tillers in maize^28^. Differences between the teosinte and maize alleles of *teosinte glume architecture1* abolished the hard seedcase which products teosinte seeds^29^. The gene *grassy tillers1* has been implicated in reducing the number of ears produced by maize^8^. The classical maize gene *zfl2* maps close to a QTL controlling differences in ear rank between maize and teosinte which would be consistent with the known function of the mutant phenotype^30^. The panicoid specific gene *ramosa1* regulates inflorescence branching and was an apparent target of selection based on single gene analysis of maize and teosinte alleles^31^. The gene *early flowering4* is located in a genomic region involved in photoperiod perception in temperate inbreds and tropical inbreds which contributes to the adaptation of temperate maize lines to the altered growing seasons and photoperiods of higher latitudes^32^. ZmCCT is another gene which maps to a QTL controlling flowering time at temperate latitudes and mutations of this gene have been show to reduce photoperiod sensitivity in tropical maize lines^33^. Genes involved in starch biosynthesis – including *amylose extender1, brittle endosperm2, sugary1, starch synthase1,* and *starch branching enzyme3* also show evidence of selection in maize relative to teosinte, which is consistent with the changes in seed composition seen during domestication and improvement^34,35^. *Opaque2 heterodimerizing proteins2* is a transcription factor which has been shown to regulate production of the storage protein zein^36^. The circadian clock-associated gene *gigantea* suppresses flowering in long day photoperiods^37^. Allelic variation at *barren inflorescence2* in maize effects tassel architecture traits such as plant height, node number, leaf length, and flowering time^38^. *zea agamous2* as the target of artificial selection which is associated with development of female inflorescence^39^. *Gln synthetase* was identified as a gene that contributes to increase nitrate in leaves under low nitrogen condition^40^. The *colorless2* gene included in maysin related QTL which was required for high silk maysin synthesis to resistant to corn earworm^41^.

**Table 1.** Well Characterized Domestication Genes in Maize and Their Orthologs/Homeologs.

| Gene symbol | Gene name | Gene ID in sorghum | Gene ID in maize1 | Gene ID in maize2 |
| --- | --- | --- | --- | --- |
| sh1 | Seed shattering1 | Sobic.001G152901 | GRMZM2G085873 | GRMZM2G074124 |
| gt1 | Grassy tillers1 | Sobic.001G468400 | GRMZM2G005624 | NoGene |
| tga1 | Teosinte glume architecture1 | Sobic.007G193500 | NoGene | GRMZM2G101511 |
| tb1 | Teosinte branched1 | Sobic.001G121600 | AC233950.1_FG002 | AC190734.2_FG003 |
| ra1 | Ramosa1 | Sobic.002G197700 | GRMZM2G003927 | NoGene |
| ELF4 | Early flowering 4 | Sobic.002G193000 | GRMZM2G025646 | NoGene |
| CCT | Flowering time | Sobic.002G275100 | GRMZM2G179024 | GRMZM2G367834 |
| ohp2 | Opaque2 zein storage protein synthesis | Sobic.001G056700 | GRMZM2G007063 | NoGene |
| yab14 | Yabby14 | Sobic.006G160800 | GRMZM2G054795 | GRMZM2G005353 |
| GI | GIGANTEA | Sobic.003G040900 | GRMZM5G844170 | GRMZM2G107101 |
| zag2 | Zea agamous2 | Sobic.008G072900 | GRMZM2G010669 | GRMZM2G160687 |
| bif2 | Barren inflorescence2 | Sobic.008G170500 | GRMZM2G171822 | NoGene |
| zfl2 | Zea floricaula/leafy2 | Sobic.006G201600 | GRMZM2G180190 | GRMZM2G098813 |
| gln2 | glutamine synthetase2 | Sobic.001G116400 | GRMZM2G024104 | NoGene |
| sbe3 | starch branching enzyme3 | Sobic.006G066800 | GRMZM2G073054 | GRMZM2G128395 |
| c2 | colorless2 | Sobic.005G136200 | GRMZM2G422750 | GRMZM2G151227 |

Note: Genes in red, cyan, and black represent the genes identified as likely under selection during domestication or improvement phases, or not showing evidence of selection in either process, respectively.

### Population genetic datasets for both species

Maize and sorghum accessions were sampled from published datasets^14,42,43^ (See Methods and Table S1). After quality filtering to remove low quality SNPs and those potentially representing alignments of paralogous sequence elsewhere in the genome (see Methods), a total of 10.3M segregating SNPs in 56 maize accessions and 3.3M segregating SNPs in 42 sorghum accessions remained. These proportions roughly correspond to the difference in genome size between the two species – approximately 2.0 gigabases for maize and 700 megabases for sorghum – however, as much of the maize genome is repetitive and cannot be uniquely identified using short sequence reads, this is consistent with higher overall levels of nucleotide diversity in maize relative to sorghum. Also consistent with previous reports, wild relatives had higher levels of nucleotide diversity (*π* =0.00377 in maize and *π* =0.00381 in sorghum), than both landraces (*π* =0.00338 in maize and *π* =0.00242 in sorghum) and improved inbreds (*π* =0.00334 in maize and *π* =0.00226 in sorghum) (Table S3)^12,14^.

Maize accessions were primarily collected in the western hemisphere and sorghum accessions primarily from the eastern hemisphere with some exceptions in both cases (Fig 1A). Wild relatives were primarily collected near the centers of domestication: southwestern Mexico for maize and central east Africa for sorghum. The sorghum dataset also included data for a forage sudangrass line (Greenleaf: sweet sorghum x Sudan grass) from North America. Among the maize lines, modern elite lines and wild relatives each formed monophyletic groups (Fig 1B). In sorghum, wild relatives formed a monophyletic clade but lines reported to be elite or landrace lines were intercalated, potentially as a result of distinct sorghum breeding efforts developing lines for different agroclimactic zones around the world (Fig 1C).

**Figure 1.**
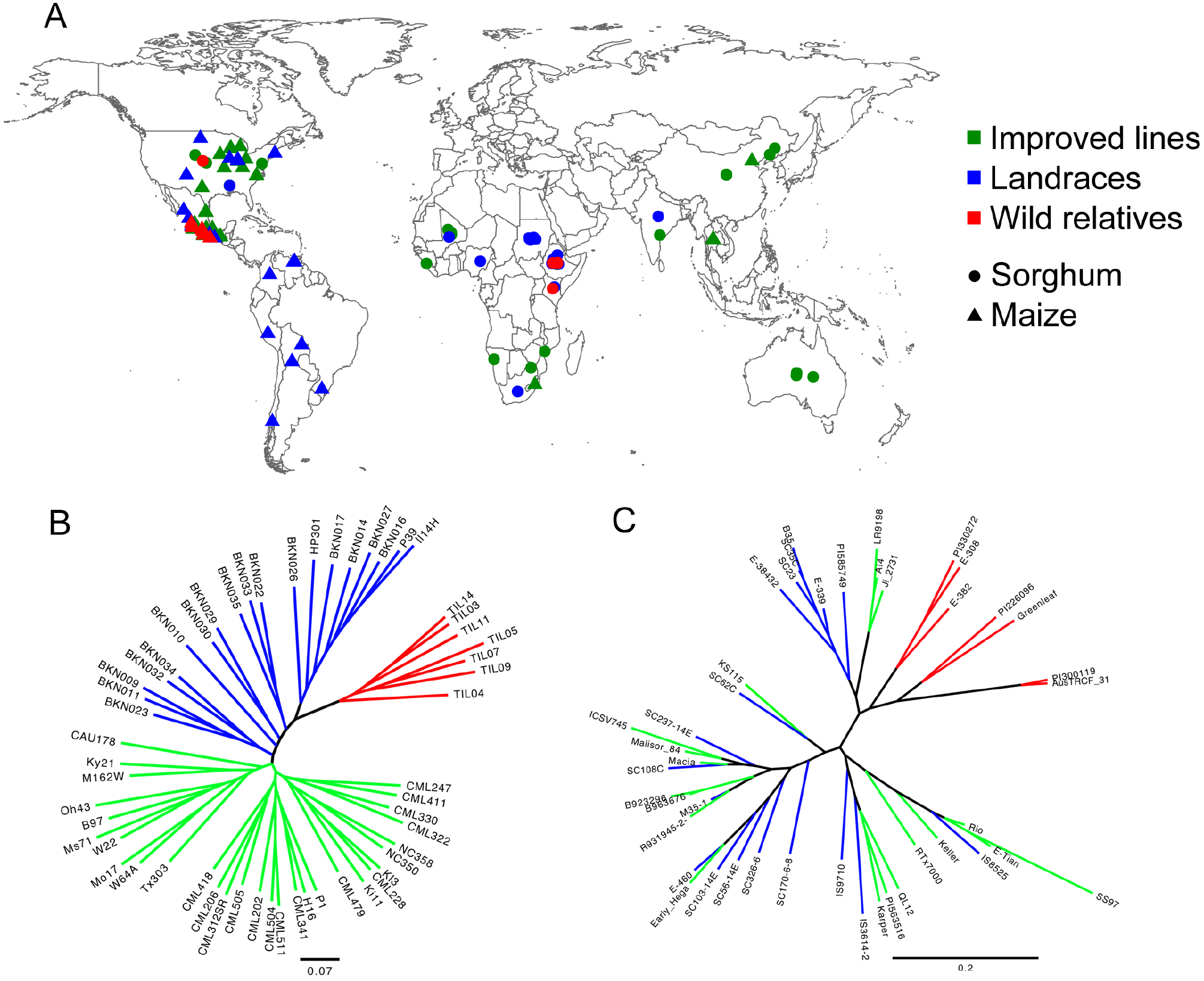
Geographic distribution and phylogenetic relationship of maize and sorghum accessions employed in this study. A) The geographic distribution of maize and sorghum accessions. Dots and triangles represent sorghum and maize, respectively. Colors represent membership in different populations: improved lines (green), landraces (blue), and wild relatives (red). B-C) Neighbor-joining trees for maize accessions (B) and sorghum accessions (C). Taxa in the NJ tree are color coded using the same system described for subpanel A.

Genetic maps for both species were sourced from public datasets. For maize, a genetic map was employed that included 10,085 markers genotyped in a set of 232 RILs from the maize IBM population using tGBS^44^ while for sorghum a genetic map was employed which was constructed from a set of 3,418 markers genotyped in a set of 244 RILs from a grain sorghum x sweet sorghum cross using resequencing^45^.

### Genomic signals of selection in maize and sorghum

In each species, the identification of regions under selection performed for three separate pairwise comparisons: landraces versus wild relatives (domestication), improved lines versus landraces (improvement), and improved lines versus wild relatives. The genome was scanned in 1kb bins using XP-CLR (see Methods) and each gene was assigned the XP-CLR score of the highest 1 kb bin overlapping with the gene. Genes above the 90th percentile of XP-CLR scores for a given pairwise comparison were considered as candidate “under selection” genes. The set of gene annotations employed in this analysis included 63,480 maize gene models and 34,027 sorghum gene models. Thus, for each of the three possible pairwise comparisons, 6,348 genes in maize and 3,403 genes in sorghum were identified as candidates for selection (Figure S1). In both species, estimated selection coefficients were higher during domestication (mean s = 0.06 in maize and 0.047 in sorghum) than improvement (mean s = 0.045 in maize and 0.024 in sorghum).

Since 10% of genes were identified as candidates for selection during domestication and 10% of genes were identified as candidates for selection during improvement, the overlap expected if these datasets were unrelated is 1%. In fact, approximately 1% of all annotated maize genes (620 genes) were in the top 10% in both comparisons and approximately 1% of all annotated sorghum genes (345 genes) were in the top 10% in both comparisons (Figure 2A-B). As expected, the set of candidate genes identified in the comparison of wild relatives and improved lines showed significant overlap with both the domestication and crop improvement candidate gene sets (Figure 2A). Among a set of 112 genes identified and cloned by classical maize using forward genetics^46^, 23 were identified as candidate genes under selection during domestication and/or improvement (p-value = 0007, Fisher exact test).

**Figure 2.**
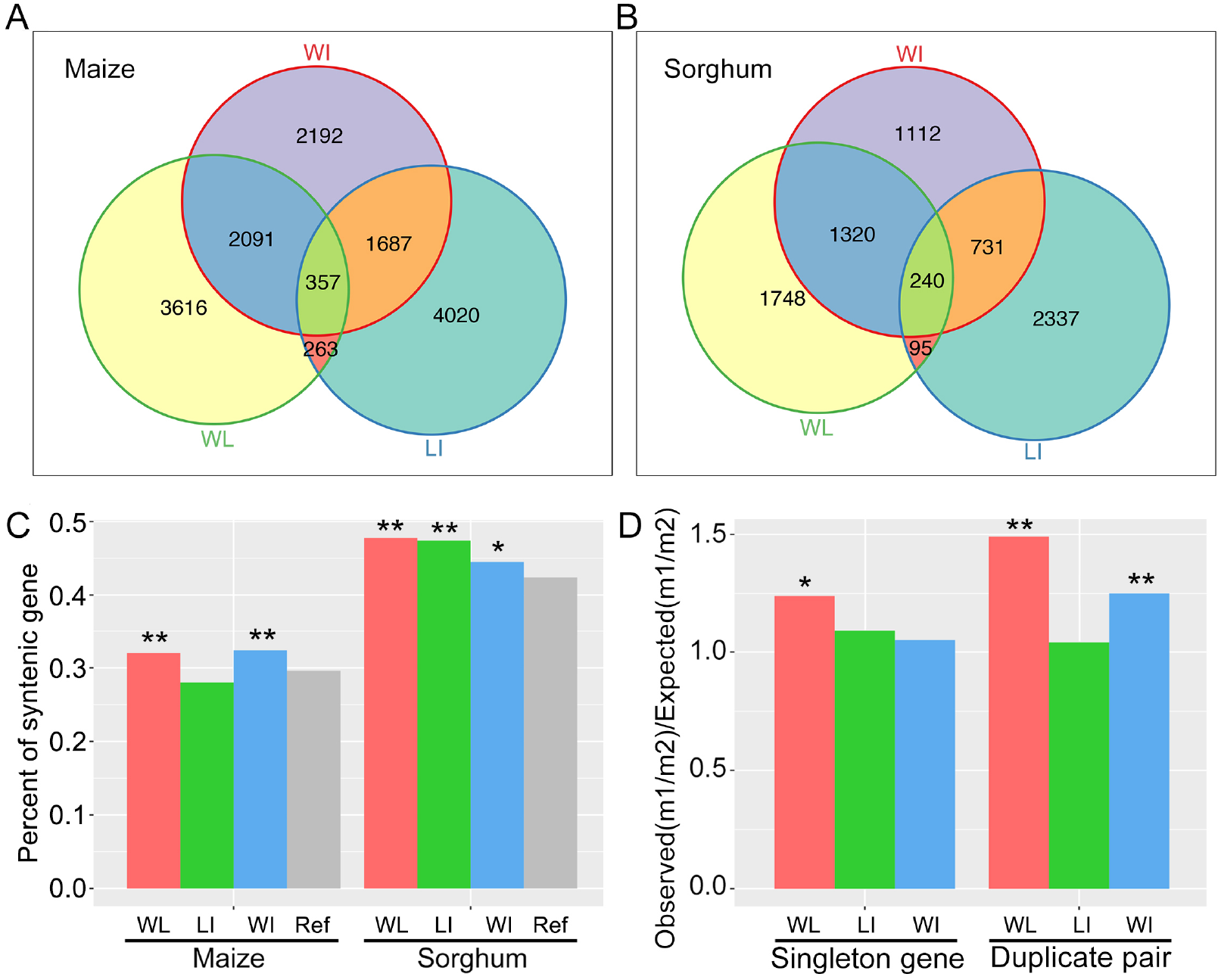
Summary information for candidate genes under selection. A-B) The number of candidate genes shared among the three contrasts in maize (A) and sorghum (B), respectively. C) The proportion of non-syntenic genes and syntenic genes under selection in pairwise comparison in maize and sorghum. D) Ratio of maize1:maize2 genes among genes identified as selection candidates. Analysis conducted separately for singleton genes and duplicate genes. One asterisk denotes cases which are significantly different at a threshold of p <0.01 and two asterisks denote cases which were significantly different at a threshold of p <0.001. I: improved lines, L: landraces, and W: wild relatives.

The maize and sorghum genetic maps are largely collinear^26^, however maize has experienced a whole genome duplication relative to sorghum, producing two functionally distinct subgenomes with differing levels of gene expression, purifying selection, and gene loss rates^47–49^. In a dataset of 14,433 genes conserved at syntenic locations in sorghum and maize 7,041 genes conserved between sorghum and maize1 but lost from maize2 and 3,031 genes conserved between sorghum and maize2 but lost from maize1, while the balance – 4,361 genes – are present in all three genomes (Table S2)^50^. The balance of the gene complement of each species consists of nonsyntenic genes, which are the majority (57.6% and 70.4% for sorghum and maize respectively) of all annotated genes in both species. Genes with known mutant phenotypes tend to be syntenic^46^, while nonsyntenic genes tend to exhibit greater allelic variation in gene regulation, and it has been speculated that they may contribute to phenotypic plasticity and heterosis^51–53^.

In sorghum, genes conserved at syntenic orthologous locations between maize and sorghum were significantly more likely to be identified as targets of selection in all three pairwise comparisons (binomial testing p-value < 1e-6 for the domestication and improvement, p-value = 0.0058 for the wild-improved comparison). In maize, syntenic genes were significantly more likely to be identified as targets of selection during domestication and in the the wild relative improved line comparison (p-value = 0.00024 and p-value = 1.0e-6 respectively) however genes identified as targets of selection during the crop improvement process were significantly more likely to be nonsyntenic genes (p-value = 0.0026) (Figure S3C).

In maize, selection candidates were also unevenly distributed between subgenomes. Maize1 genes were more likely to be identified as candidates for selection during domestication both among gene retailed as duplicate pairs (1.49x) and genes which fractionated to single copy (1.24x), (p-value = 0.00013 and p-value ¡ 1e-6 respectively, binomial test) (Figure 2D). Fewer singleton genes were identified as likely to be under selection during domestication than duplicate gene pairs while the opposite pattern was observed for genes identified as likely to be under selection during improvement, however, these differences were not statistically significant (Figure S2).

### Testing for parallel selection during domestication

In order to control for differences in gene content and biases towards syntenic genes, new sets of candidate genes was selected consisting of only genes above the 90th percentile for XP-CLR scores of syntenically conserved genes in each species. Based on the percentage of genes classified as domestication candidates in each species, in the absence of parallel selection on the same genes during domestication 189 gene pairs would be expected to be identified as gene candidates in both species. Among domestication candidate genes 196 gene pairs were identified independently in both species, slightly more than the 189 gene pairs expected in the absence of parallel selection (determined via permutation testing). This difference of 7 genes was not statistically significant (FDR ¡ 0.27, permutations) (Figure 3A). The gene exhibiting the strongest combined selection signal across the two species GRMZM2G026024/Sobic.004G272100 encodes a phosphoribulokinase, an enzyme that catalyzes a key step in carbon fixation as part of the Calvin cycle.

**Figure 3.**
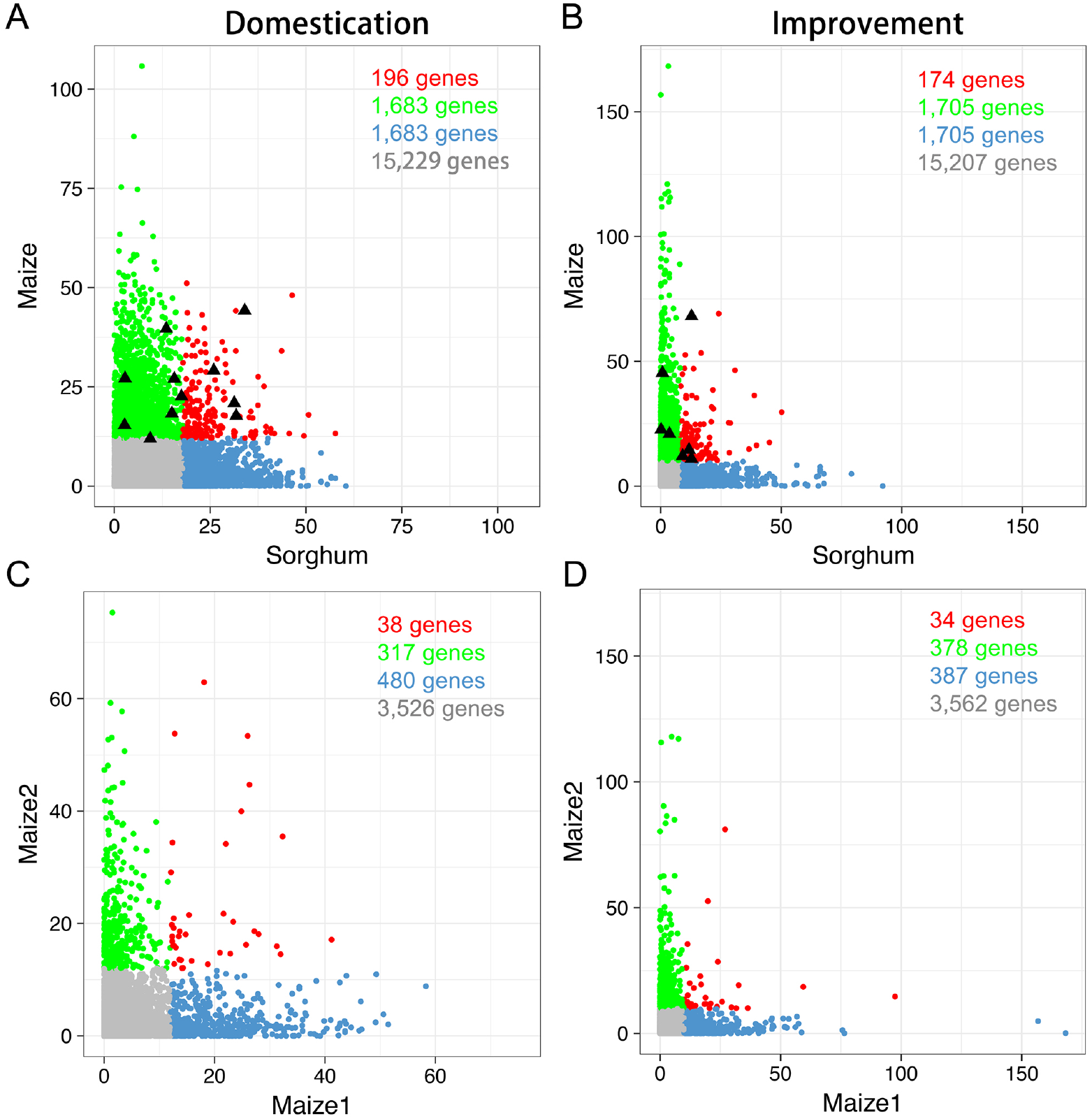
Comparison of scores for syntenic orthologous gene pairs in the wild relatives/landrance (A) and landrace/improved lines XP-CLR analyses of maize and sorghum as well as the selection in duplicated maize genes during the domestication (C) and improvement (D). Red, blue, orange, and black dots (in A and B) mark gene pairs identified as putative selection candidates in both maize and sorghum, only in sorghum, only in maize, or in neither species, respectively. The triangles in A and B show the classic domestication genes of maize listed in table 1. And red, blue, orange, and black dots (in C and D) mark gene pairs identified as putative selection candidates in both maize1 and maize2, only in maize1, only in maize2, or in neither subgenome, respectively.

Comparison of genes under apparent selection in the landrace vs elite comparison identified fewer pairs syntenic genes under parallel selection than during the domestication (Figure 3B). This may be linked to a lower proportion of the syntenic genes under selection in maize during the improvement process (Figure 2C). In the case of genes under selection during crop improvement a total of 186 overlapping genes were expected but 174 were observed (FDR ¡ 0.85, permutations) (Figure 3B). In the wild relatives vs elite comparison 195 overlapping gene pairs were identified and 188 were expected (FDR ¡ 0.31, permutations) (Figure S3C).

The cut off of genes in the 90th percentile of XP-CLR scores was chosen somewhat arbitrarily. In order to test whether the lack of greater than expected overlap between genes identified in maize and those identified in sorghum was an artifact of the threshold score employed the analysis above was repeated using a range of percentile based score thresholds from the 85th percentile to the 99th percentile. None of these thresholds identified a significant enrichment of gene pairs under selection in both species relative to the expectations of the null hypothesis (Figure S4A,B).

Another potential explanation is that the analysis above was partially confounded as a result of the partially paired data structure, with some sorghum genes paired with a single maize syntenic ortholog and others paired with two syntenic orthologs on opposite maize subgenomes. Separate permutation tests were conducted using only maize1-sorghum or maize2-sorghum gene pairs. Slightly more gene pairs were identified as likely under selection during improvement between maize1 and sorghum than expected under the null hypothesis and slightly fewer gene pairs than expected were identified in the maize2 sorghum comparison. However, this bias was not large and was not replicated in the comparison of genes identified as likely under selection during domestication (Figure S4C-F).

While many phenotypic changes during domestication appear to be shared between sorghum and maize – the domestication syndrome referenced above – domestication likely also involved selection on some traits only in one species or the other. Therefore, we also searched for signatures of parallel selection between homeologous gene pairs retained between the two subgenomes of maize, as these genes experienced identical whole-plant level artificial selection during the domestication of maize. 38 duplicate pairs under selection during domestication were identified from 4,362 pairs of retained maize duplicates tested (Figure 3C) and only 34 duplicate pairs under parallel selection during improvement were identified (Figure 3D). Smaller number of duplicate genes under parallel selection during the improvement because of a smaller proportion of syntenic genes under selection during this period. In both cases, the number of gene pairs identified as likely under parallel selection were lower than the expectation for unlinked genes, but not by a statistically significant amount (FDR ¡ 0.88 and 0.80, permutations, respectively).

Another potential explanation for the absence of significant overlap between genes appearing to have experienced selection in maize and sorghum, or between the two maize subgenomes, is simply that the dataset used or analysis method employed was invalid in some way. To test this concern, employed the positive control set of 16 maize genes with known and functionally validated links to domestication phenotypes in maize described above (Table 1). All sixteen of these genes were indeed included in the set of maize gene candidates identified through XP-CLR analysis. In nine cases the sorghum orthologs of these target genes were also identified as likely targets of artificial selection (p-value < 1.0e-06, binomial test).

Characterized genes showing signatures of parallel selection in both maize and sorghum include the previously reported *sh1* gene involved in the loss of seed shattering^15^. Genes involved in reshaping plant architecture such as *gt1,* identified as a controller of ear number in maize^8^, and genes involved in regulation of flowering time such as *ELF4* and GI^54^, as did two important genes in starch synthesis pathway *ss1* and sbe3^34,55^. *tb1* which is involved in the repression of axillary branching in both maize^28^ and sorghum^56^ showed signatures of parallel selection, which was identified as parallel selection candidate in maize and sorghum. However, the *tb*1 was not one of the strongest signals of selection even was not found to be candidate when included the *mexicana* lines^12^. A second TCP transcription factor, belonging to the same gene family as *tb1,* was identified as under parallel selection in sorghum, maize1, and maize2 (Figure 4).

**Figure 4.**
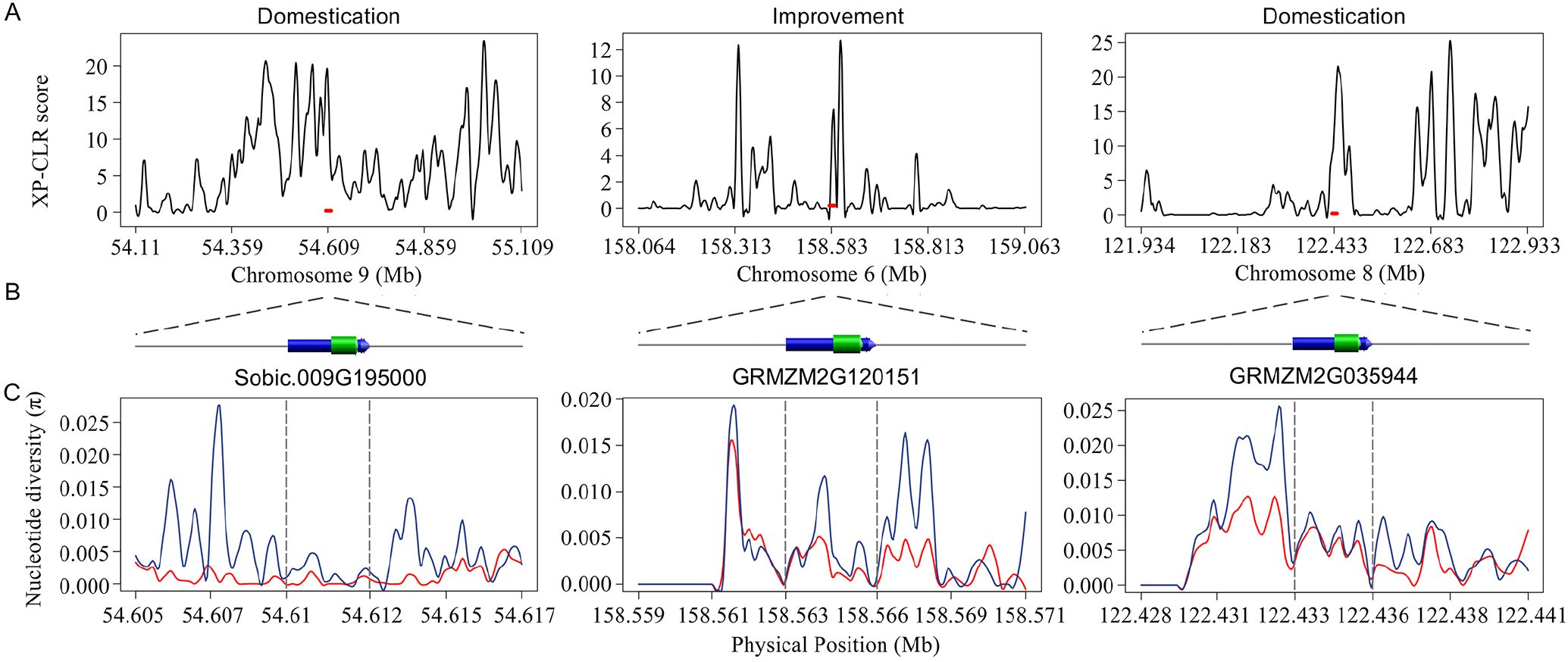
An example of evidence of selection on a *TCP* gene in maize and sorghum. A) Cross-population composite likelihood ratio test plot showing selection on partial regions of genomes in sorghum, maize1, and maize2. Red line represents the location of orthologous *TCP* gene on the chromosomes. B) Gene model of syntenic *TCP* genes in sorghum, maize1, and maize2. The blue and green boxes represent the UTR regions and Exons. C) Level of nucleotide diversity (*π*) in objective population (red) and background population (blue).

### Functional roles of genes selected in parallel

A total of 1,014 maize/sorghum syntenic gene pairs were identified as under parallel selection including both genes under parallel selection in the same comparison (ie wild vs landrance or landrace vs improved) or in opposite comparisons in different species. These genes were enriched in transcription factors relative to all syntenic gene pairs in both sorghum (p-value = 8.70e-4, Fisher Exact Test) and maize (p-value = 3.50e-5, Fisher Exact Test), although the absolute enrichment is modest (1.36x and 1.46x for maize and sorghum respectively) (Table S4).

A total of 237 domestication candidate genes in sorghum were involved in 112 annotated biochemical pathways and 270 domestication candidate genes from the 18,794 syntenic genes of maize were involved in 113 annotated biochemical pathways. A total of 69 pathways overlapped between these two datasets, which was similar to the 71 pathways predicted to overlap based on permutations of orthologous relationships (FDR ¡ 0.606, permutations). For genes identified as candidates in the landrace-improved line comparison, a total of 231 candidate genes from sorghum were annotated as being part of 128 pathways and 258 selection candidate genes from maize were annoated as being part of 129 pathways. A total of 80 pathways had at least one maize genes and one sorghum genes under selection which is moderately statistically significant compared to the expected number of overlapping pathways (FDR ¡ 0.006, permutations). However, when analyzing gene pairs identified as under parallel selection, a set of 141 gene pairs were annotated as encoding enzymes which were involved in 89 different metabolic pathways (Table S4). This was a significantly larger number of metabolic pathways than would be expected given the overall number of gene pairs under parallel selection (Expectation = 78 pathways, FDR ¡ 0.028, permutations).

A set of 44 maize sorghum gene pairs identified as selection candidates in both species were also found to exhibit tissue specific expression RNA-seq data generated from homologous tissues in maize and sorghum^57,58^, particularly in anthers, embryos, and endosperm (Table S5). One gene which showed strong parallel selective signals in sorghum (Sobic.009G203900) and both maize subgenomes – maize1 (GRMZM2G074361) and maize2 (GRMZM2G109842) – exhibited identical and highly specific expression patterns in the anthers of both species (Figure S5). This gene is annotated profilin 1 (PRF1) gene which encodes a core cell-wall structural proteins. Previous studies have reported that PRF1 can mediate cell signaling and in crop plants can alter flowering time and the number of floral meristems produced by inflorescence meristems^59,60^.

## Discussion

The high collinearity of genetic maps and gene content among related grasses^26^ makes it feasible to employ multiple grass species as a single genomic system. Here we sought to test whether the parallel phenotypic changes produced by artificial selection as part of the domestication syndrome in maize and sorghum resulted from parallel molecular changes targeting orthologous genes. However, as shown above, the number of genes showing parallel signatures of selection during domestication in maize and sorghum was not significantly different from the amount of overlap expected among random gene sets. This result stands in contrast to reports from individual large effect genes where the same gene often appears to have been a target during independent domestication in different species^15,16^, as well as the finding here that genes with validated links to domestication from single gene studies in maize where significantly more likely to also be targets of selection in sorghum (Table 1).

A recently report which compared published results from analyses of domestication in maize and rice reached a similar result with only 65 orthologous gene pairs shared between a set 969 genes identified as candidate targets of selection in maize and 1,526 gene pairs identified as candidate targets of selection in rice^61^. However, that study also identified a number of limitations of their analysis, including the relatively high LD in rice, and the potential for selection for different traits during domestication in the two species given the large differences in growth habit between modern rice and maize cultivars, and that the analysis reused candidate gene sets identified in different research groups using different sets of parameters. Here we employed data from two more closely related species with similar – and low – levels of LD and greater similarities of plant architecture and growth habit as well as conducting a reanalysis starting from raw SNP calls in order to ensure balanced and equivalent approaches to identifying candidate genes in both species. However, as reported by Gaut, we also find an absence of parallel selection on orthologous genes at a whole genome level between maize and sorghum.

We have included a number of control analysis to validate this result. A set of positive control domesticate candidate genes were used to ensure that the statistical methods, software implementations, and genomic datasets employed do indeed have the power to identify genes which were targets of selection during domestication and crop improvement. Analysis of parallel selection between the subgenomes of maize was used to rule out the explanation that the lack of overlap between maize and sorghum orthologs resulted from selection for different traits during domestication. Analysis of biochemical pathways identified no evidence that artificial selection in maize and sorghum tended to targeted different genes in the same pathways. The absence of signatures of parallel selection could be explained by gain-of-function mutations targeting different family members in different species for co-option into the same new functional roles, as appears to have happened with the SH1 gene in rice and the AP2 gene in wheat^62^. It may also be that the same genes have different levels of importance in regulating the same traits in different species. For example the *tb1* gene which has such a large effect on tillering in maize has been show to have a much smaller role in controlling tillering in foxtail millet^63^. Another potential explanation is that domestication largely acts on preexisting standing variation within a wild progenitor species and which genes were targets of artificial selection is largely determined by which alleles were segregating in the target wild population. Genes such as *tb1, zfl2, zagl1*, and *zap1* all show evidence of exhibiting standing functional variation in teosinte prior to the initial domestication of maize^12,64,65^. However, it must be noted that there are indeed significant differences in the domestication process between the two species. Maize was domesticated from teosinte approximately 9,000 years ago in what appears to have been a single event^21,66^. Improved maize lines used in this study were largely drawn from temperate elite varieties adapted to North America. In contrast, sorghum appears to have been domesticated at least twice independently^14^, and the improved sorghum lines used here were drawn from separate breeding efforts aimed at developing improved cultivars for African, Australian, and North American climates. In addition, selection for higher yield in maize has resulted in indirect selection for decreases in tassel size^67^, while selection for decreased head size in sorghum would presumably be detrimental to yield.

The observation that the few maize domestication genes characterized through single gene conventional genetic studies were much more likely to also be identified as domestication candidates in sorghum (Table 1) does suggest that the genes involved in domestication may fall into a two-tier system. A few large effect genes appear to have been repeatedly targeted to create the domestication syndrome in multiple grain crops^15,16^. As described by Orr, strong and relatively recent selection should rapidly identify and fix a small number of large effect alleles which may have pleiotropic consequences, while a a range of different smaller effect alleles at other genetic loci are then selected to fine tune the effect size and mitigate any negative pleiotropic effects of the initial large effect alleles^68^. Studies of the genetic architecture of different traits inferred to be under selection or largely neutral characters during domestication and crop improvement in maize, and in maize-teosinte RIL populations have produced findings consistent with this model^69,70^. Here we propose that the initial, large effect alleles selected for during selection for domestication syndrome traits in grain crops are drawn from a constrained pool of genes, and therefore orthologous genes are more likely to be selected for in parallel across multiple grain crops. In contrast the set of small effect genes which fine tune domestication rates and mitigate potentially deleterious pleiotropic effects of large effect alleles, may drawn from a much larger pool and would thus exhibit little repeat sampling of the same orthologous genes across different domesticated grasses. The decreasing cost of whole genome sequencing and resequencing, and the large number of different grain crops which have experienced parallel selection for domestication syndrome phenotypes should enable more rigorous tests of this model in the near future incorporating data from syntenic orthologous genes across many different species.

## Materials and Methods

### Data collection and preliminary polishing

Maize and sorghum whole genome resequencing data used in this study were taken from Hapmap2^42^ and SorGSD^43^ respectively. SNPs scored as heterozygous in ≥ 3.0% of accessions in maize hapmap 2 and sorghum dataset were removed prior to analysis.

A subset of maize accessions selected and separated into three groups: 30 improved lines, 19 landraces, and 7 wild relatives (Table S1). A total of 42 sorghum accessions having a spanning dimensions of geographic origin were obtained, including 17 improved lines, 18 landraces, and 7 wild relatives (Table S1).

SNPs with missing rates >50% in either species, or with heterozygous calls in any of the remaining samples were discarded, resulting in a final dataset consisted of 10.3 million SNPs in maize and 3.3 million SNPs in sorghum.

### Population genetics analysis

The genetic distance between individuals was first calculated using a 0.1% subset of total SNP set constructed by sampling every 1,000th SNP position along each chromosome for a total of 10,286 SNPs in maize and using a 0.2% subset of the total SNP set constructed by sampling every 500th SNP position along each chromosome for a total of 6,719 SNPs in sorghum.

Neighbor-joining trees were constructed for the accessions of each species using Phase^71^ and Phylip v3.696^72^ with default parameters. The resulting phylogenetic trees were visualized using Figtree v1.4.3 (http://tree.bio.ed.ac.uk/software/figtree/). Nucleotide diversity (*π*) values were calculated for each species with non-overlapping windows of 10 kb using an in-house Perl script. Reported *π* values are the average of all genomic windows.

### Syntenic gene identification

A pan-grass syntenic gene set using the sorghum genes as reference were download from the figshare^50^. When multiple sorghum genes identified as syntenic orthologs of the same gene in maize – a result which can be produced by tandem duplication events in sorghum – the tie was broken using a separate dataset of syntenic orthologous genes using the *Setaria italica* genome as a reference. This resulted in a final set of 14,433 sorghum genes paired with a syntenic ortholog in either the maize1 subgenome (11,402 gene pairs) and/or the maize2 subgenome (7,392 gene pairs), including 4,361 sorghum genes with syntenic co-orthologs on both maize subgenomes (Table S2).

### Genome-wide scan for selection

To identify genes affected by selection during the domestication in maize and sorghum, genome-wide scans for signals of selection were conducted using a cross-population composite likelihood approach (XP-CLR)^73^ (updated by Hufford et al.^12^ to incorporate missing data), based on the allele frequency differentiation between target and reference populations. This approach was employed in three separate pairwise comparisons: wild relatives vs landraces, landraces vs improved lines, and wild relatives vs improved lines.

Recombination rates in maize and sorghum were measured using high density genetic maps in maize^44^ and sorghum^45^. Genetic maps were transfered to the more recent versions of the maize and sorghum genomes used in this analysis (B73_RefGen_v3 and v3.1 respectively). The transfer was performed using the two genes flanking the marker – when the marker was in a noncoding region – or the single gene the marker was located in. For each pseudomolecule in each species, a ninth order polynomial curve was fit to the genetic and physical coordinates of all markers presented on chromosomes, and genetic positions for each marker were reassigned based on the value predicted for genetic and physical position of the marker and the polynomial formula.

The same parameters were employed for XP-CLR analysis for maize and sorghum. A 0.05 cM sliding window with 1,000 bp steps across the whole genome scan was used for scanning. Individual SNPs were assigned a position along the genetic map by assuming uniform recombination between pairs of genetic markers. Equivalent results were obtained when estimated SNP genetic positions using the polynomial formula estimated above (data not shown). The number of SNPs assayed in each window was fixed at 100 and pairs of SNPs in high LD (r2 > 0.75) were down-weighted.

To obtain XP-CLR scores for each gene, each gene was assigned a window starting 5 kb upstream of its annotated transcription start site and extending to 5 kb downstream of its annotated transcription stop site. The maximum XL-CLR score among all the XL-CLR intervals within this window was assigned to the gene.

### Testing for Enrichment of Genes Selected in Parallel

Gene pairs were considered to be under selection if a sorghum gene and at least one maize syntenic ortholog were both identified as being under selection. To determine the optimal cut-off for testing the enrichment of syntenic genes under parallel selection, a series of cut-offs from 85% to 99% were used in the analysis. At each cut off the number of gene pairs under selection in both species were recorded and compared to the number of gene pairs identified when orthologous relationships between maize and sorghum were shuffled using a permutation test repeated for 100 times.

### Gene annotation and enrichment analysis

Maize and sorghum GO annotations were retrieved from phytozome (https://phytozome.jgi.doe.gov/). The maize transcription factor (TF) lists were downloaded from Grassius (http://grassius.org/grasstfdb.html). Metabolic pathway lists were download from the Gramene (ftp://ftp.gramene.org/pub/gramene/pathways/). Annotated enzyme name and the correspond pathways for these genes were obtained by searching the pathway list.

A set of 1,000 permutations were used to calculate the expected number of pathways in the same number of random genes in R package to examine whether maize and sorghum genes under selection were significantly more likely to be present in the same pathways than expected if selection was unlinked.

## Acknowledgements

We thank Prof. Ed Buckler (Cornell University) for advice on the hapmap dataset and Prof. Jeffrey Ross-Ibarra (UC Davis) for advice an access to an updated version of the XP-CLR software. This work was supported by a China Scholarship Council fellowship awarded to XL and a Science Foundation of Xichang College awarded to LY.

## Author contributions

JCS and XL conceived the project and designed the studies; XL and LY performed the research; XL, LY, JCS, and YL wrote the paper. All authors reviewed the manuscript.

## Figure legends

**Figure 1.** Geographic distribution and phylogenetic relationship of maize and sorghum accessions. A) The geographic distribution of maize and sorghum accessions. Dots and triangles represent sorghum and maize, respectively. Colored squares represent the populations: improved lines (green), landraces (blue), and wild relatives (red). B-C) Neighbor-joining tree of maize (B) and sorghum (C). Taxa in the NJ tree are represented by different colors: improved lines (green), landraces (blue), and wild relatives (red).

**Figure 2.** Summary information of candidate gene under selection. A-B) The number of candidate genes shared among the three contrasts in maize (A) and sorghum (B), respectively. C) Proportion of non-syntenic genes and syntenic genes under selection in pairwise comparison in maize and sorghum. D) Ratio of maize1:maize2 genes among genes identified as selection candidates. Analysis conducted separately for singleton genes and duplicate genes. One asterisk denotes the significant level P <0.01 and double denote the significant level P <0.001. I: improved lines, L: landraces, and W: wild relatives.

**Figure 3.** Comparison of scores for syntenic orthologous gene pairs in the wild relatives/landrance (A) and landrace/improved lines XP-CLR analyses of maize and sorghum as well as the selection in duplicated maize genes during the domestication (C) and improvement (D). Red, blue, orange, and black dots (in A and B) mark gene pairs identified as putative selection candidates in both maize and sorghum, only in sorghum, only in maize, or in neither species, respectively. The triangles in A and B show the classic domestication genes of maize listed in table 1. And red, blue, orange, and black dots (in C and D) mark gene pairs identified as putative selection candidates in both maize1 and maize2, only in maize1, only in maize2, or in neither subgenome, respectively.

**Figure 4.** An example of evidence of selection on *TCP* genes in maize and sorghum. A) Cross-population composite likelihood ratio test plot showing selection on partial regions of genomes in sorghum, maize1, and maize2. Red line represents the location of orthologous *TCP* gene on the chromosomes. B) Gene model of syntenic *TCP* genes in sorghum, maize1, and maize2. The blue and green boxes represent the UTR regions and Exons. C) Level of nucleotide diversity (*π*) in objective population (red) and background population (blue).

## Supplemental Information (SI)

**Table S1** List of maize and sorghum accessions employed in this study, their data sources, and their classifications as wild relative, landrace, or improved line datasets.

**Table S2** Set of high confidence syntenic orthologous maize sorghum gene pairs employed in this study.

**Table S3** Population nucleotide diversity statistics for maize and sorghum.

**Table S4** A list of the genes identified as likely under selection during the domestication and/or improvement process in maize and sorghum and functional annotations of each.

**Table S5** Tissue-specific expression genes under parallel selection between maize and sorghum.

**Figure S1** Genome-wide scan for signatures of selection in maize and sorghum. Each dot represents the XP-CLR score assigned to a single gene.

**Figure S2** Ratio of singleton genes:duplicate pair among genes identified as selection candidates. I: improved lines, L: landraces, and W: wild relatives.

**Figure S3** Comparison of scores for syntenic orthologous gene pairs in the wild relative/improved line XP-CLR analyses of maize and sorghum. Red, blue, orange, and black dots mark gene pairs identified as putative selection candidates in both maize and sorghum, only in sorghum, only in maize, or in neither species, respectively. The triangles show the classic domestication genes of maize listed in table 1.

**Figure S4** Testing for enrichment of candidate selection genes for different proportion of genes with top XP-CLR score.

**Figure S5** Gene model and gene expression level visualization of *profilin1* in sorghum and maize. A) Gene model visualization of syntenic gene *profilin1* in Coge (https://genomevolution.org/CoGe/GEvo.pl). B-D) Gene expression level of *profilin1* in different tissues of sorghum (B), maize1 (C), and maize2 (D) obtained from qteller (www.qteller.com).

**Figure S1.**
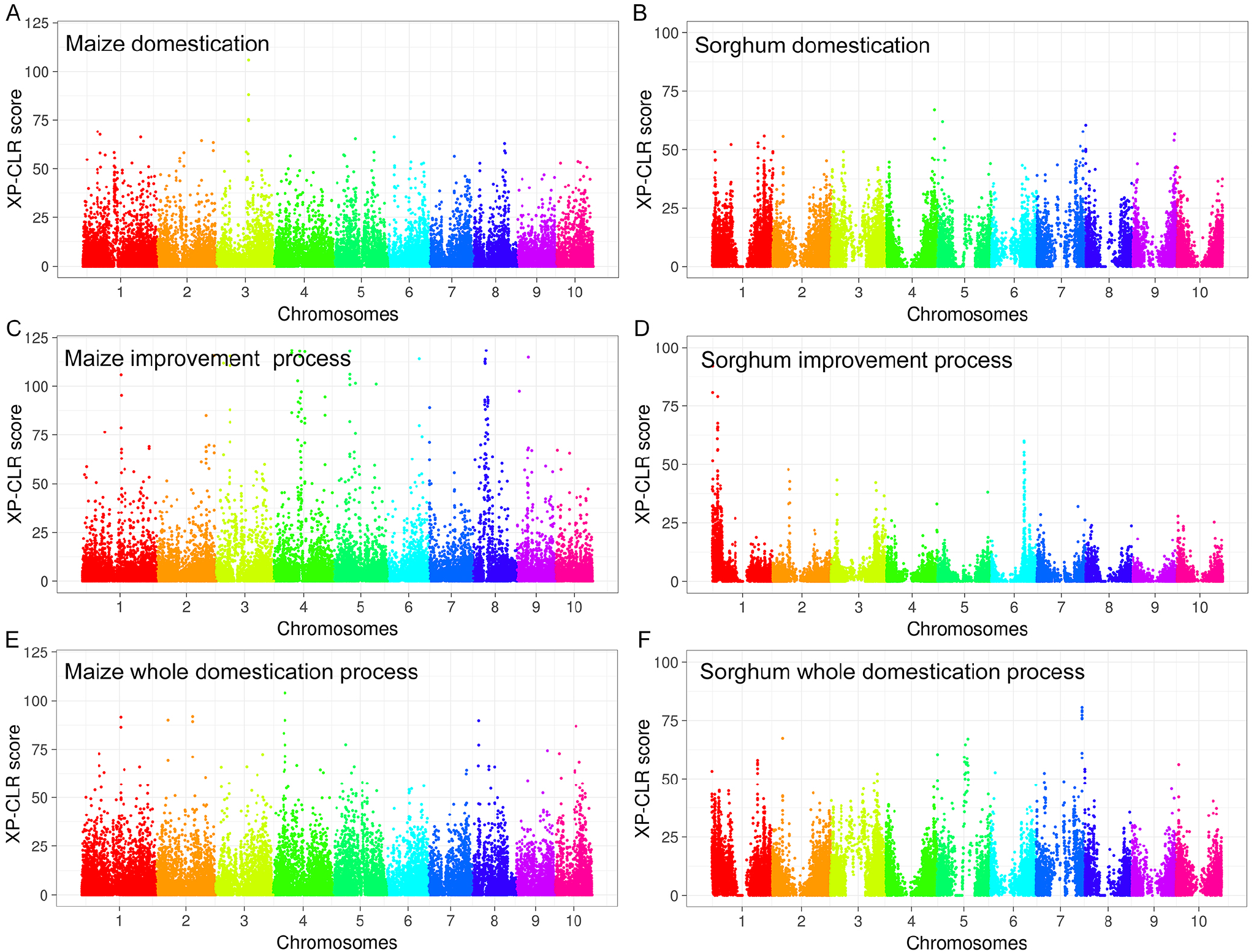
Genome-wide scan for signatures of selection in maize and sorghum. Each dot represents the XP-CLR score assigned to a single gene.

**Figure S2.**
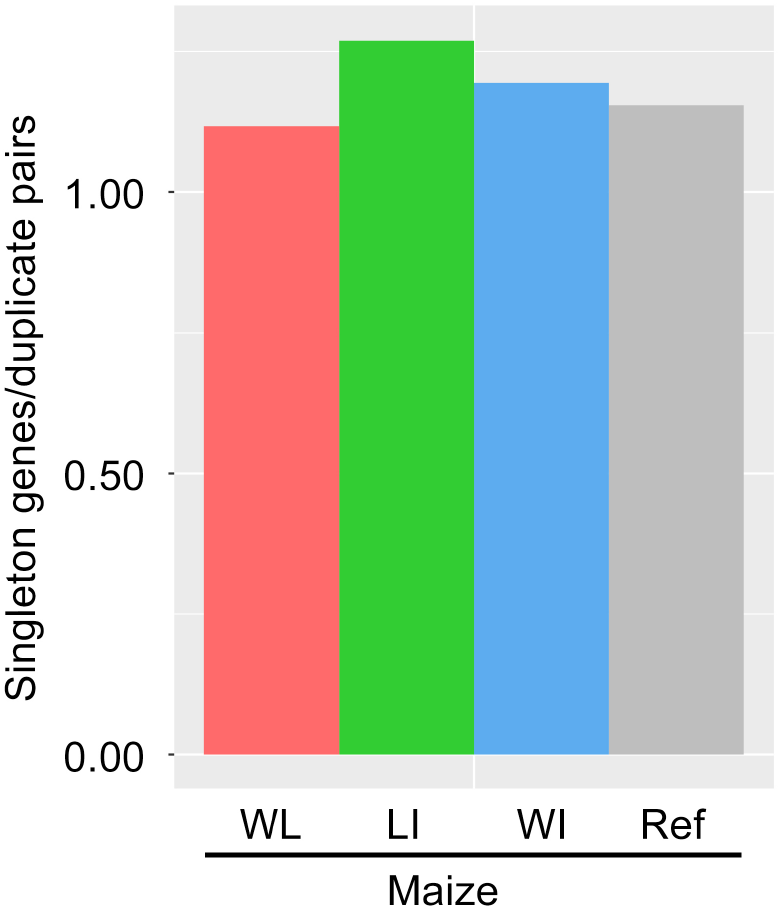
Ratio of singleton genes:duplicate pair among genes identified as selection candidates. I: improved lines, L: landraces, and W: wild relatives.

**Figure S3.**
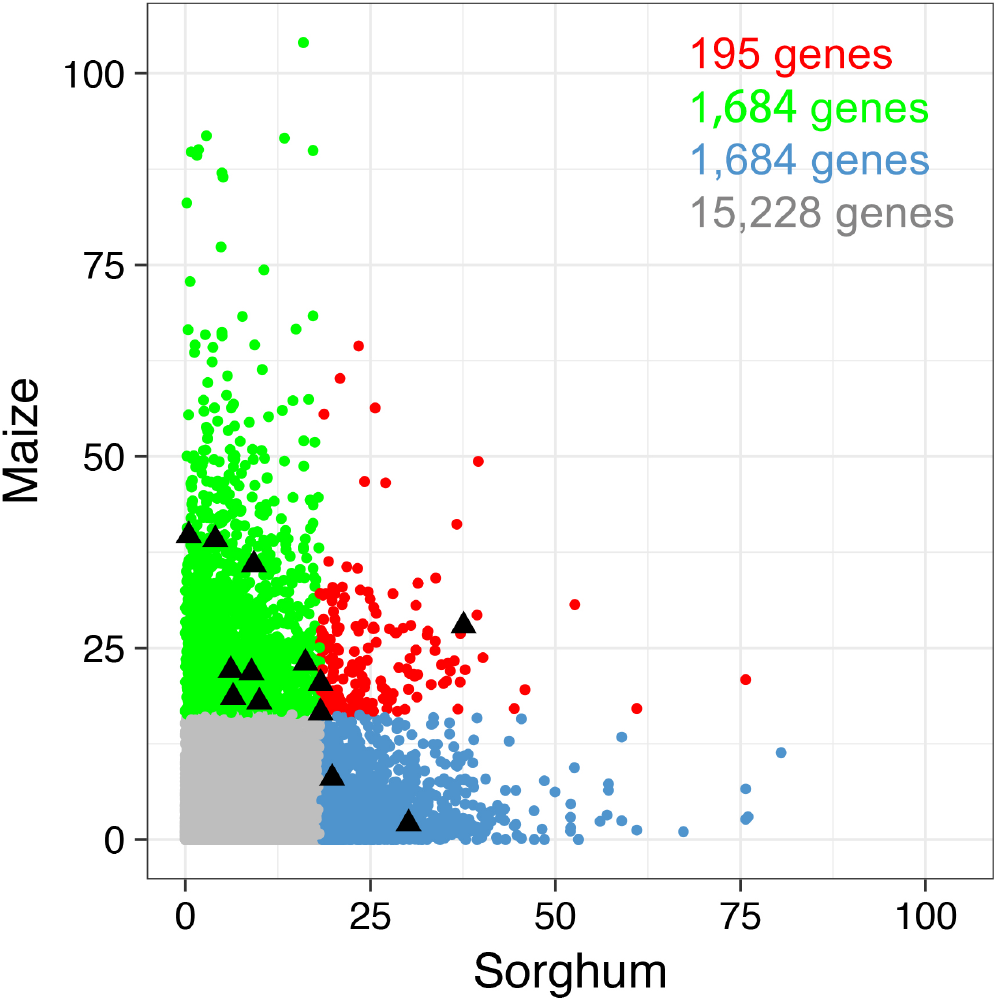
Comparison of scores for syntenic orthologous gene pairs in the wild relative/improved line XP-CLR analyses of maize and sorghum. Red, blue, orange, and black dots mark gene pairs identified as putative selection candidates in both maize and sorghum, only in sorghum, only in maize, or in neither species, respectively. The triangles show the classic domestication genes of maize listed in table 1.

**Figure S4.**
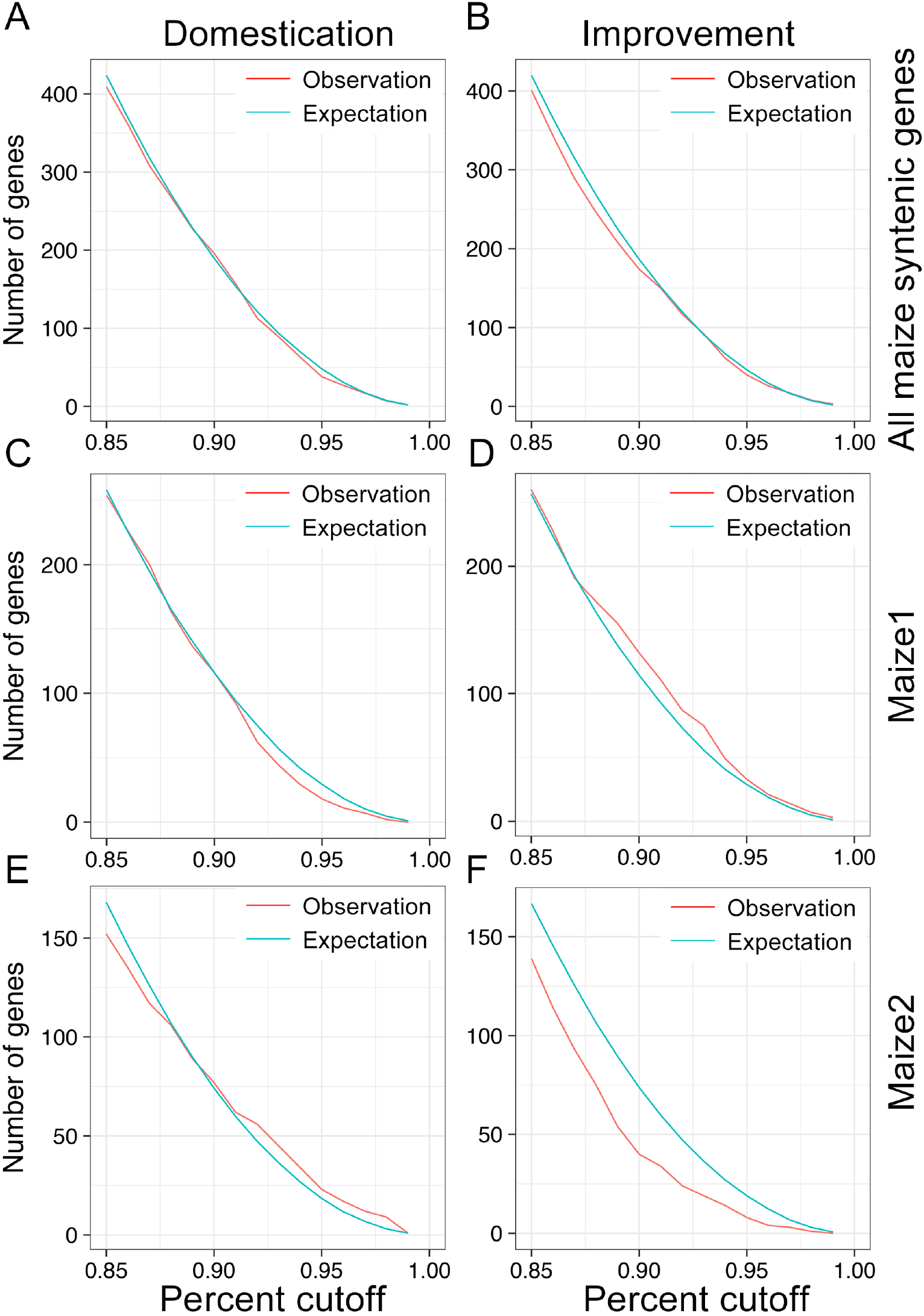
Testing for enrichment of candidate selection genes for different proportion of genes with top XP-CLR score.

**Figure S5.**
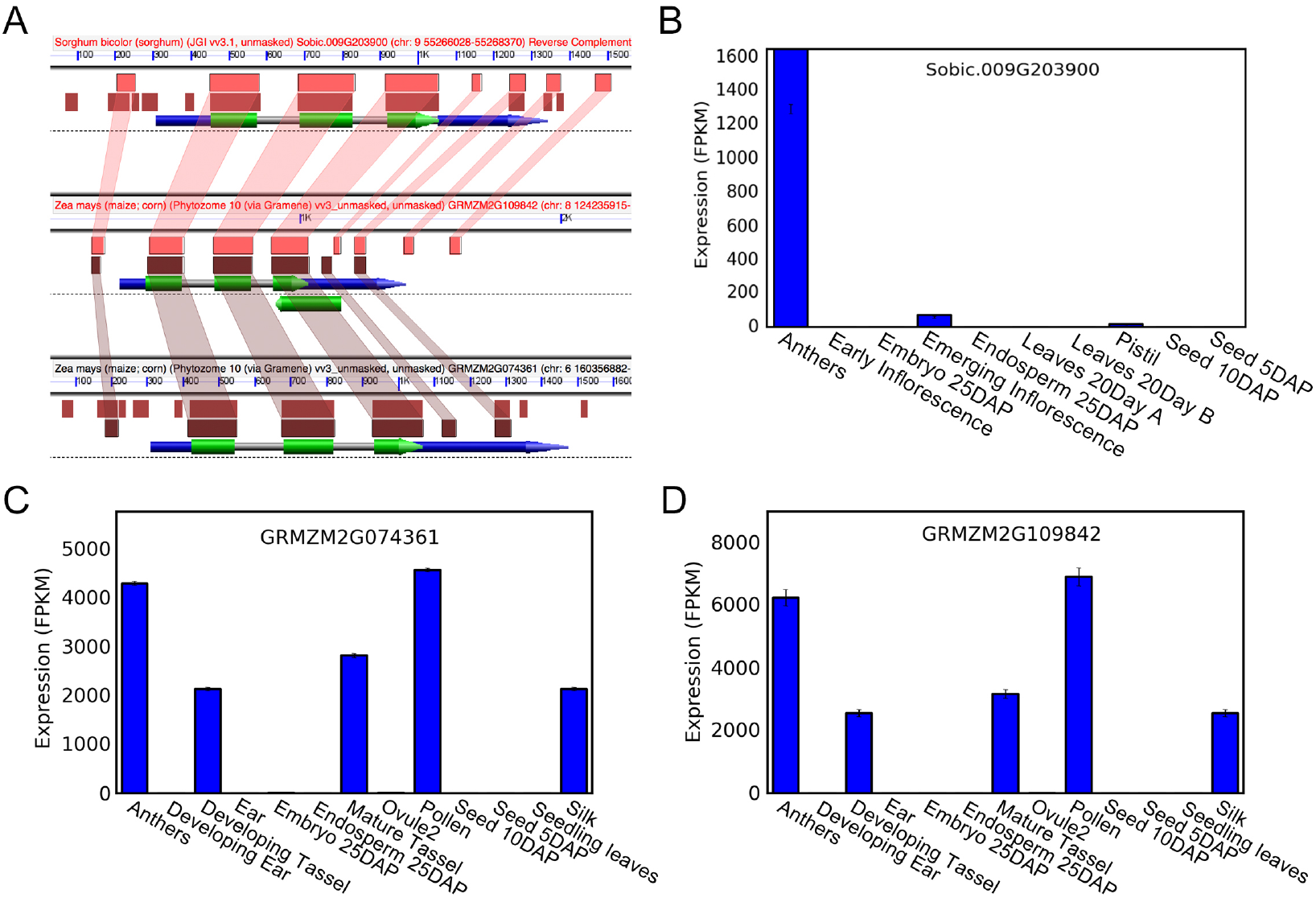
Gene model and gene expression level visualization of *profilin1* in sorghum and maize. A) Gene model visualization of syntenic gene *profilinl* in Coge (https://genomevolution.org/CoGe/GEvo.pl). Gene expression level of *profilin1* in different tissues of sorghum (B), maize1 (C), and maize2 (D) obtained from qteller (www.qteller.com).

